# Two RNA binding proteins, ADAD2 and RNF17, interact to form novel meiotic germ cell granules required for male fertility

**DOI:** 10.1101/2022.11.11.516086

**Authors:** Lauren G. Chukrallah, Sarah Potgieter, Elizabeth M. Snyder

## Abstract

Mammalian male germ cell differentiation relies on complex RNA biogenesis events, many of which occur in RNA binding protein (RBP) rich non-membrane bound organelles termed RNA germ cell granules. Though known to be required for male germ cell differentiation, little is understood of the relationships between and functions of the numerous granule subtypes. ADAD2, a testis specific RBP, is required for normal male fertility and forms a poorly characterized granule in meiotic male germ cells. This work aimed to define the role of ADAD2 granules in male germ cell differentiation and their relationship to other granules. Biochemical analyses identified RNF17, a testis specific RBP that forms meiotic male germ cell granules, as an ADAD2-interacting protein. Phenotypic analysis of *Adad2* and *Rnf17* mutant mice defined a shared and rare post-meiotic chromatin defect, suggesting shared biological roles. We further demonstrated ADAD2 and RNF17 are dependent on one another for granularization and together form a previously unstudied set of germ cell granules. Based on co-localization studies with well-characterized granule RBPs including DDX4 and PIWIL1, a subset of the ADAD2-RNF17 granules are likely components of the piRNA pathway. In contrast, a second, morphologically distinct population of ADAD2-RNF17 co-localize with the translation regulator NANOS1 and form a unique cup-shaped structure with distinct protein subdomains. This cup shape appears to be driven, in part, by association with the endoplasmic reticulum. Lastly, a double *Adad2-Rnf17* mutant model demonstrated loss of ADAD2-RNF17 granules themselves, as opposed to loss of either ADAD2 or RNF17, is the likely driver of the *Adad2* and *Rnf17* mutant phenotypes. Together, this work identified a set of novel germ cell granules required for male fertility and sheds light on the relationship between germ cell granule pools. The example described here defines a new genetic approach to germ cell granule study.

**AUTHOR SUMMARY:** To differentiate successfully, male germ cells tightly regulate their RNA pools. As such, they rely on RNA binding proteins, which often localize to cytoplasmic granules. The majority of studies have focused on a single granule type which regulates small-RNA biogenesis. Although additional granules have been identified, there is limited knowledge about their relationship to each other and exact functions. Here, we identify an interaction between two RNA binding proteins, ADAD2 and RNF17, and demonstrate mutants share a rare germ cell phenotype. Further, ADAD2 and RNF17 colocalize to the same germ cell granule, which displays two morphologically unique types. The first subset of ADAD2-RNF17 granules have similar morphologies to other characterized granules and likely play a role in the small-RNA pathway. The second granule type forms a unique shape with distinct protein subdomains. This second population appears to be closely associated with the endoplasmic reticulum. Genetic models further demonstrate the granules themselves, as opposed to the resident proteins, likely drive the mutant phenotypes. These findings not only identify a novel population of germ cell granules but reveal a new genetic approach to defining their formation and function during germ cell differentiation.

## INTRODUCTION

The male germ cell relies on complex RNA biology for successful differentiation. As a result, they express and require a wide range of RNA binding proteins (RBPs). Many RBPs are housed in non-membrane bound, cytoplasmic organelles termed germ cell RNA granules or germ cell granules. These granules are especially prevalent during meiotic and post-meiotic germ cell differentiation and are fundamental for proper developmental progression in multiple species [1,2]. In mammalian male germ cells for example, six types of granules have been identified via electron microscopy (EM), five of which can be found in meiotic spermatocytes [3]. Loss of core granule proteins commonly leads to meiotic or post-meiotic germ cell arrest [4–8], underscoring their importance in germ cell differentiation and male fertility.

Historically, the function of these granules has been defined primarily by functional knowledge of associated proteins. One particularly successful example of this is the intermitochondrial cement (IMC), which is an amorphous matrix distributed between the mitochondria of meiotic male germ cells (spermatocytes) with well-defined protein and RNA composition [6,9–11]. Based on a combination of genetic and functional analyses, the IMC has been identified as a primary site of biogenesis for piRNAs [7,9,12–14], an abundant class of small non-coding RNAs that modulate mRNA translation and stability [15,16]. Many IMC-localized proteins, including the primary piRNA biogenesis factors PIWIL1 [17] and PIWIL2 [18] have distinct impacts on the piRNA biogenesis events localized to the IMC [7,14].

Unlike the IMC, the other meiotic germ cell granules have less defined compositions or functions. Three (the satellite or sponge body, loose aggregate strands, and irregularly shaped perinuclear granules) are known to contain RBPs associated with mRNA storage and translation regulation such as DDX4, DDX25, and NANOS1 [19–21]. In even greater contrast to the IMC is the one entirely unstudied meiotic germ cell granule, known only as the “cluster of 30-nm particles”, which has no defined resident proteins and has only been observed via electron microscopy. In addition to relatively poor composition and functional knowledge for the non-IMC granules, the exact functional role of or relationship between them has never been defined [22]. Similarly, whether the cluster of 30-nm particles or any of the other EM-defined granule types represent single, molecularly homogenous populations has been almost entirely unexplored. Together these questions represent a long-standing mystery in male germ cell RNA biology.

One promising approach to address these mysteries involves detailed studies of RBPs that form spermatocyte germ cell granules. Of particular interest are those proteins that have yet to be assigned to a specific granule population. ADAD2, a testis specific RBP [23], forms a spermatocyte germ cell granule and is required for successful male germ cell development as *Adad2* mutant males are completely infertile, with germ cell development halting abruptly during post-meiotic germ cell differentiation [23]. Appearing first in pachytene spermatocytes wherein it is largely cytoplasmic, ADAD2 coalesces into a distinct perinuclear granule during mid-meiosis and remains thus through the end of meiosis. Little is known about the composition of the ADAD2 granule beyond its lack of DDX25 [23], known to mark all the spermatocyte germ cell granules excluding the cluster of 30-nm particles [20,24].

To define the ADAD2 granule, we set out to identify additional protein components of the ADAD2 granule and relate the granule’s composition to other meiotic granules. Using the mouse as a model and leveraging multiple single and complex genetic models as well as high-resolution imaging modalities, we further dissect potential drivers of ADAD2 granule formation as well as the role of the individual proteins relative to the granule itself. Together, these studies describe novel meiotic germ cell granules that may play a unique role in the complex RNA biology of the germ cell.

## RESULTS

### ADAD2 interacts with RNF17, a testis-specific RNA binding protein

The ADAD2 granule appears to be distinct from the best characterized spermatocyte granule, intermitochondrial cement (IMC) [23]. To better determine the molecular nature of the ADAD2 granule, we immunoprecipitated ADAD2 from wildtype (n = 3) and *Adad2* mutant (*Adad2^M/M^*, n = 1) testes at 42 days post-partum (dpp) followed by mass spectrometry (IP-MS) to identify potential ADAD2-granule associated proteins (Fig 1A). Hits identified in all wildtype samples but not the mutant included well characterized post-meiotic germ cell proteins along with several RNA binding proteins (Fig 1B). Of the significant peptides identified in wildtype only, ADAD2-derived peptides represented nearly a tenth, confirming efficacy of pulldown. However, the highest number of peptides identified belong to another RNA binding protein, ring finger protein 17 (RNF17). RNF17 peptides comprised over a fifth of those identified as significant. RNF17, like ADAD2, is testis-specific and has been reported to form a spermatocyte granule [8].

**Figure 1.**
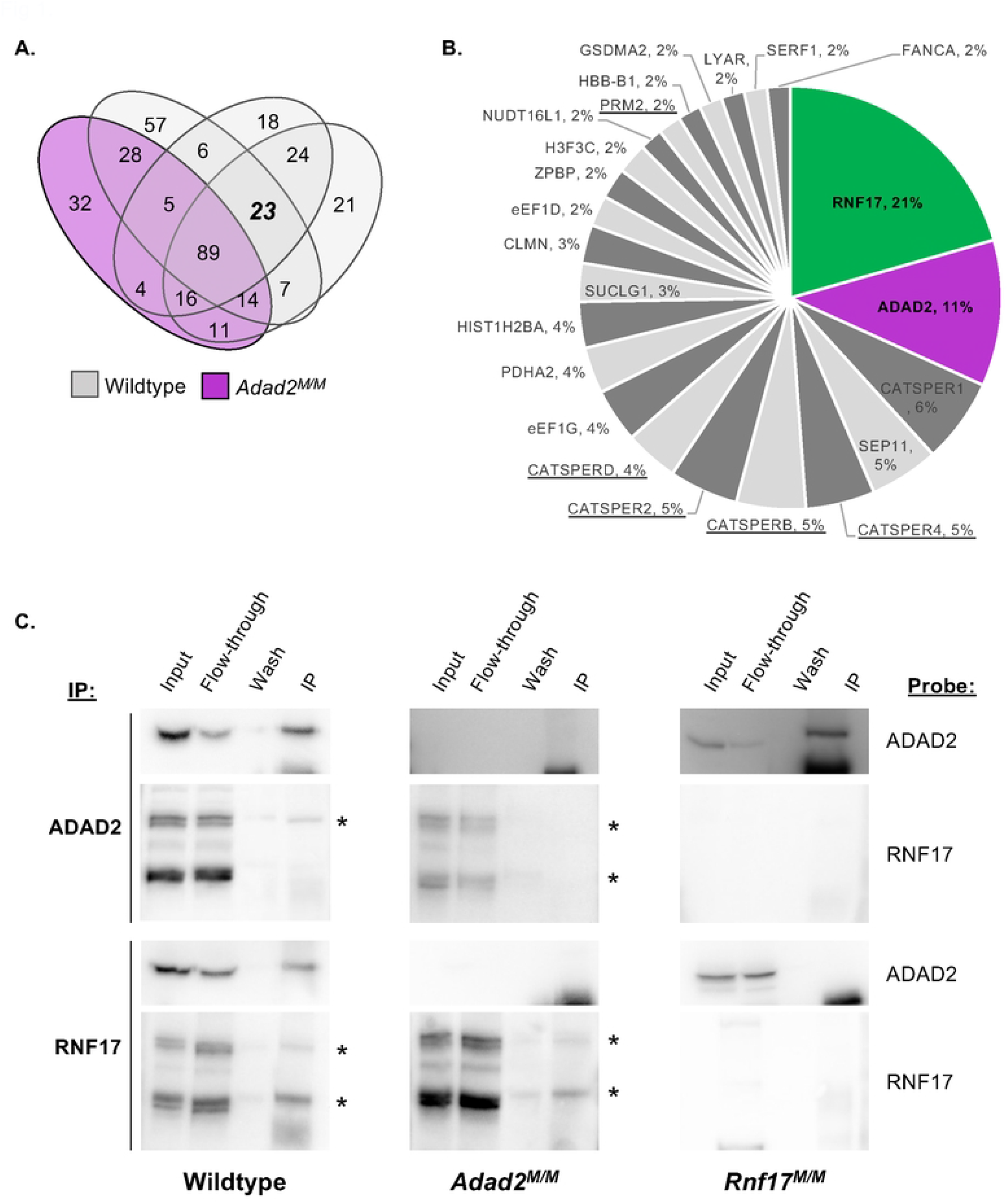
ADAD2 interacts with another RNA binding protein, RNF17. **A**. Number of mass spectrometry-identified proteins from ADAD2 immunoprecipitation (IP) of whole testis lysate from 42 dpp wildtype (grey ovals, n= 3) or *Adad2*^*M/M*^ (purple oval, n= 1). **B**. Summary of proteins detected across all wildtype IPs but not *Adad2*^*M/M*^. RNF17, in green, and ADAD2, for confirmation of genotype and IP, in purple. Underline indicates well-known post-meiotic germ cell proteins. **C**. Confirmation of ADAD2-RNF17 interaction via immunoprecipitation-Western blotting in wildtype, *Adad2* mutant (*Adad2*^*M/M*^), and *Rnf17* mutant (*Rnf17*^*M/M*^) 42 dpp whole testis lysates demonstrating a specific interaction between ADAD2 and RNF17L. Blots representative of results from at least three IPs per genotype. Asterisks - RNF17 protein isoforms.

To confirm the ADAD2-RNF17 interaction, immunoprecipitation of either ADAD2 or RNF17 in an additional set of 42 dpp wildtype testes as well as in *Adad2* mutant [23] and *Rnf17* mutant (*Rnf17^M/M^*) [8] testes was performed. The resulting immunoprecipitates (IPs) were probed for ADAD2 and RNF17 (Fig 1C). As expected, IP of either ADAD2 or RNF17 in wildtype testes resulted in robust detection of the precipitated protein. In the case of RNF17, this includes a large and small protein isoform (RNF17L and RNF17S), both of which have been detected previously [8]. Further, mutation of either *Adad2* or *Rnf17* resulted in a complete loss of ADAD2 or RNF17 detection, respectively. As expected from the IP-MS analysis, IP of ADAD2 resulted in definitive isolation of RNF17, specifically the large isoform RNF17L, while IP of RNF17 pulled down ADAD2. Lastly, mutation of either *Adad2* or *Rnf17* abrogated IP of the other. Together, these targeted IP studies confirm the initial IP-MS results and demonstrate ADAD2 and RNF17L interact in vitro.

To assess the impact of either *Adad2* or *Rnf17* mutation on the abundance of the other, we next quantified ADAD2 and RNF17 abundance in 42 dpp wildtype and mutant testes (S1A Fig). This analysis revealed loss of ADAD2 led to a reduction but not loss of RNF17, confirming the IP-MS results were not a function of protein loss in the mutant. Similarly, RNF17 loss led to reduction but not loss of ADAD2 protein along with the appearance of a smaller ADAD2 protein isoform. Given mutation of both *Adad2* and *Rnf17* results in germ cell loss at 42 dpp, we performed a similar analysis at 21 dpp (S1B Fig), a time point at which neither model should have substantial changes in cellularity. In contrast to the observations in 42 dpp testes, loss of ADAD2 lead to a slight increase of both RNF17 isoforms while RNF17 loss had minimal impact on ADAD2 abundance. While these findings suggest ADAD2 may directly influence RNF17 by modulating protein abundance, potential ADAD2-induced changes would not impact the above observed IP-based interactions, thus confirming ADAD2 and RNF17L interact in vitro.

### Both *Rnf17* and *Adad2* mutants exhibit round spermatids with abnormal chromocenters

As ADAD2 and RNF17 interact, we next wondered whether *Rnf17* mutation mimics that of *Adad2. Rnf17* mutant males have been shown to exhibit severe post-meiotic germ cell loss culminating in total male infertility [8] and previously published analyses of *Adad2* mutant testis histology suggest a similar profile of germ cell loss during round spermatid development [23]. To determine whether the post-meiotic phenotype in *Rnf17* mutants mimicked that observed with ADAD2 loss, quantification of post-meiotic round spermatid numbers as a function of stage was performed on both models (Fig 2A). This analysis demonstrated distinct round spermatid reduction in *Rnf17* mutants similar to that observed in *Adad2* mutants. Further, these analyses revealed *Rnf17^M/M^* round spermatids exhibit abnormal heterochromatin as marked by regions of intense DAPI staining and the heterochromatin mark H3K9me3 [25] (Fig 2B). Normal round spermatids contain two distinct, but associated, regions of heterochromatin, the first composed of autosomal heterochromatin and referred to as the chromocenter and the second composed of sex chromosome heterochromatin, referred to as post-meiotic sex chromatin (PMSC) [26]. The heterochromatin defect observed in *Rnf17* mutant round spermatids was also observed in *Adad2* mutants. To determine if the chromatin ultrastructure defect in *Adad2* and *Rnf17* mutant round spermatids shared a similar profile we quantified H3K9me3 foci in wildtype, *Rnf17*^*M/M*^, and *Adad2^M/M^* round spermatids (Fig 2C). As expected, wildtype round spermatids rarely contained more than a single focus, representing a normal chromocenter associated with PMSC. However, both *Adad2* and *Rnf17* round spermatids had increased numbers of H3K9me3 foci compared to wildtype and the increase in both mutant models was similar. This effect was independent of spermatid developmental stage thus impacting the entire post-meiotic germ cell population. To date, only four [27–30] other genetic models have been reported to have similar chromocenter defects, making it unusually rare. The observation of such a rare phenotype in both *Adad2* and *Rnf17* mutants indicates they may influence similar downstream events and further suggests their interaction is biologically relevant.

**Figure 2.**
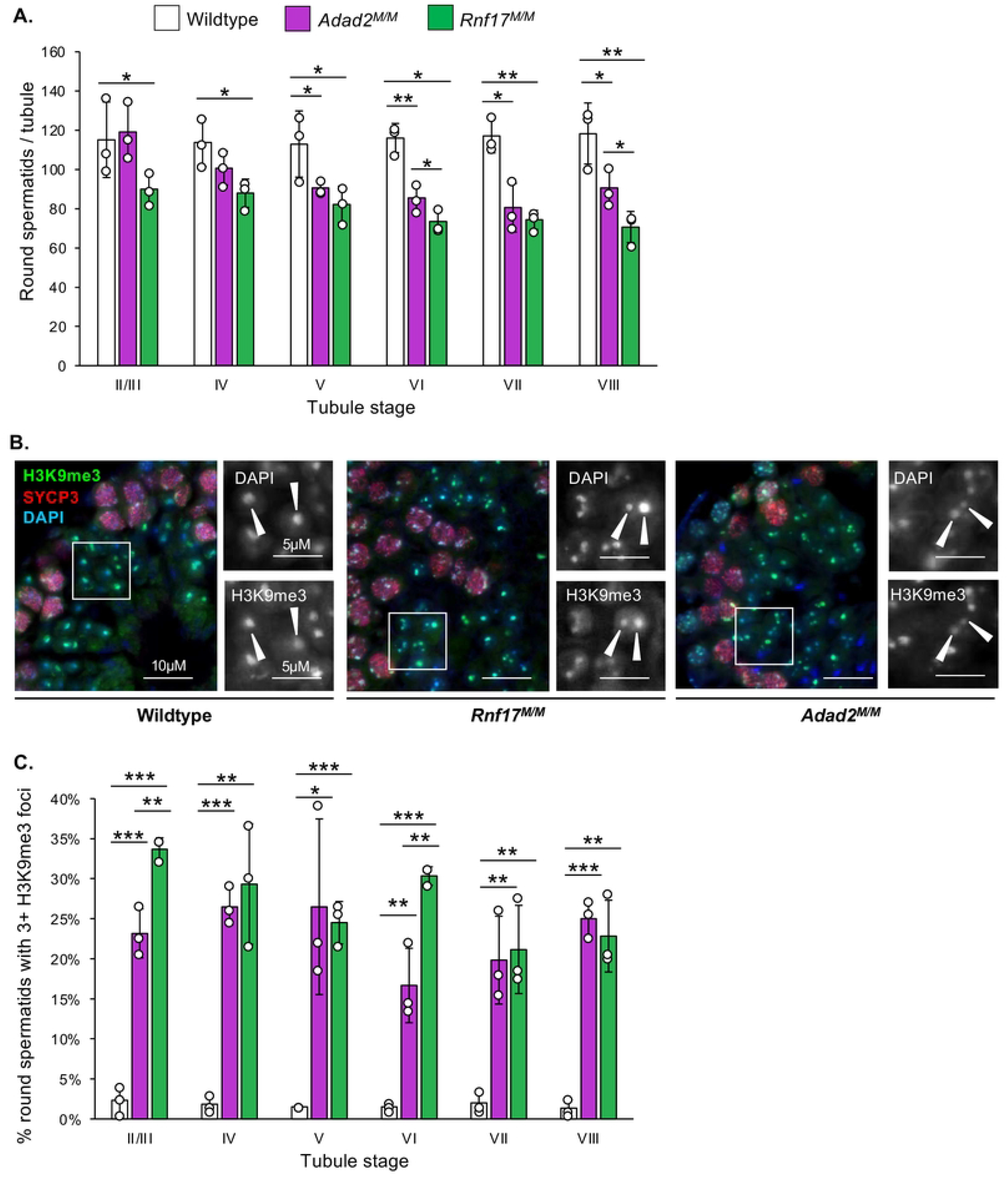
Loss of RNF17 results in a distinct chromocenter phenotype also observed in *Adad2* mutants. **A**. Round spermatids per tubule as a function of developmental stage in adult wildtype, *Adad2*^*M/M*^, and *Rnf17*^*M/M*^ testes (n = 3). **B**. Stage-matched H3K9me3 immunofluorescence in adult wildtype, *Adad2*^*M/M*^, and *Rnf17*^*M/M*^ testes counterstained with the stage-dependent marker SYCP3. Green – H3K9me3, red – SYCP3, blue – DAPI. 400x magnification. Insets – DAPI or H3K9me3 signal only. **C**. Quantification of round spermatids with 3 or more chromocenter-like structures as marked by H3K9me3 staining per developmental stage in adult wildtype, *Adad2*^*M/M*^, and *Rnf17*^*M/M*^ samples (n = 3). Data are mean ± s.d. Significance calculated using an unpaired, one-tailed Student’s t-test (\**P* < 0.05, \*\**P* < 0.005, \*\*\**P* < 0.0005).

### RNF17 has a distinct localization in spermatocytes that is dependent on ADAD2

Two RNF17 protein isoforms, large and small, have previously been reported [8]. However, IP of ADAD2 only detected RNF17L despite both isoforms being present in *Adad2*^*M/M*^ samples (see Fig 1C, S1A and S1B Figs). To determine whether ADAD2 and RNF17L shared a similar developmental profile, we examined their appearance during neonatal and juvenile development (S2A Fig). This analysis demonstrated ADAD2 first appeared at 10 dpp, coincident with the first appearance of early to mid-stage pachytene spermatocytes in the developing testis. Following this, the abundance of ADAD2 increased dramatically at 15 dpp when the testis cellular profile is highly enriched for mid-to late-pachytene spermatocytes. A very similar pattern was also observed for RNF17L. In contrast, RNF17S is observed as early as 8 dpp, reaching and sustaining a maximum by 10 dpp. Together, this demonstrates ADAD2 shares a very similar developmental profile specifically with RNF17L and further suggests ADAD2’s interaction with RNF17L, even in juvenile germ cells, would not be dependent on availability as both RNF17 protein isoforms are present from 10 dpp onward.

ADAD2 forms a developmentally-regulated granule in pachytene spermatocytes [23]. RNF17L has also been described as forming a germ cell granule in pachytene spermatocytes [8]. To better define the timing of RNF17 granule formation, we examined RNF17 localization via immunofluorescence in wildtype adult spermatocytes throughout their differentiation (Fig 3A). RNF17 weakly appears in the cytoplasm of mid-stage spermatocytes (stage V) and first coalesces into small granules in stage VII spermatocytes. Following this, the small RNF17 granules persist and additional, large cytoplasmic RNF17 granules appear between stages VIII and IX in mid-to late pachytene spermatocytes. The majority of RNF17 signal is retained in these large cytoplasmic granules until the end of meiosis (stage XII). This is similar to previous reports [8]. A similar profile is observed in juvenile spermatocytes (S2B Fig), with small and large RNF17 granules appearing at 15 dpp in mid-to late spermatocytes. Comparison with ADAD2 granule formation in the adult (S3A Fig) demonstrated ADAD2 granule formation in spermatocytes is notably delayed compared to the small RNF17 granules but aligns very well with formation of the large RNF17 granule, starting in mid-to late spermatocytes of stage VIII. Similar to RNF17, ADAD2 granules are also observed in two types, one small and frequent and the other large occurring only once or twice per cell. Across juvenile development (S2B Fig), both granule types of ADAD2 are first observed with similar timing to RNF17 granules. Together, these analyses demonstrate both ADAD2 granules and the large, but not small, RNF17 granules are specific to mid-to late pachytene spermatocytes. Further, both ADAD2 and RNF17 large granules appear at very similar developmental time points.

**Figure 3.**
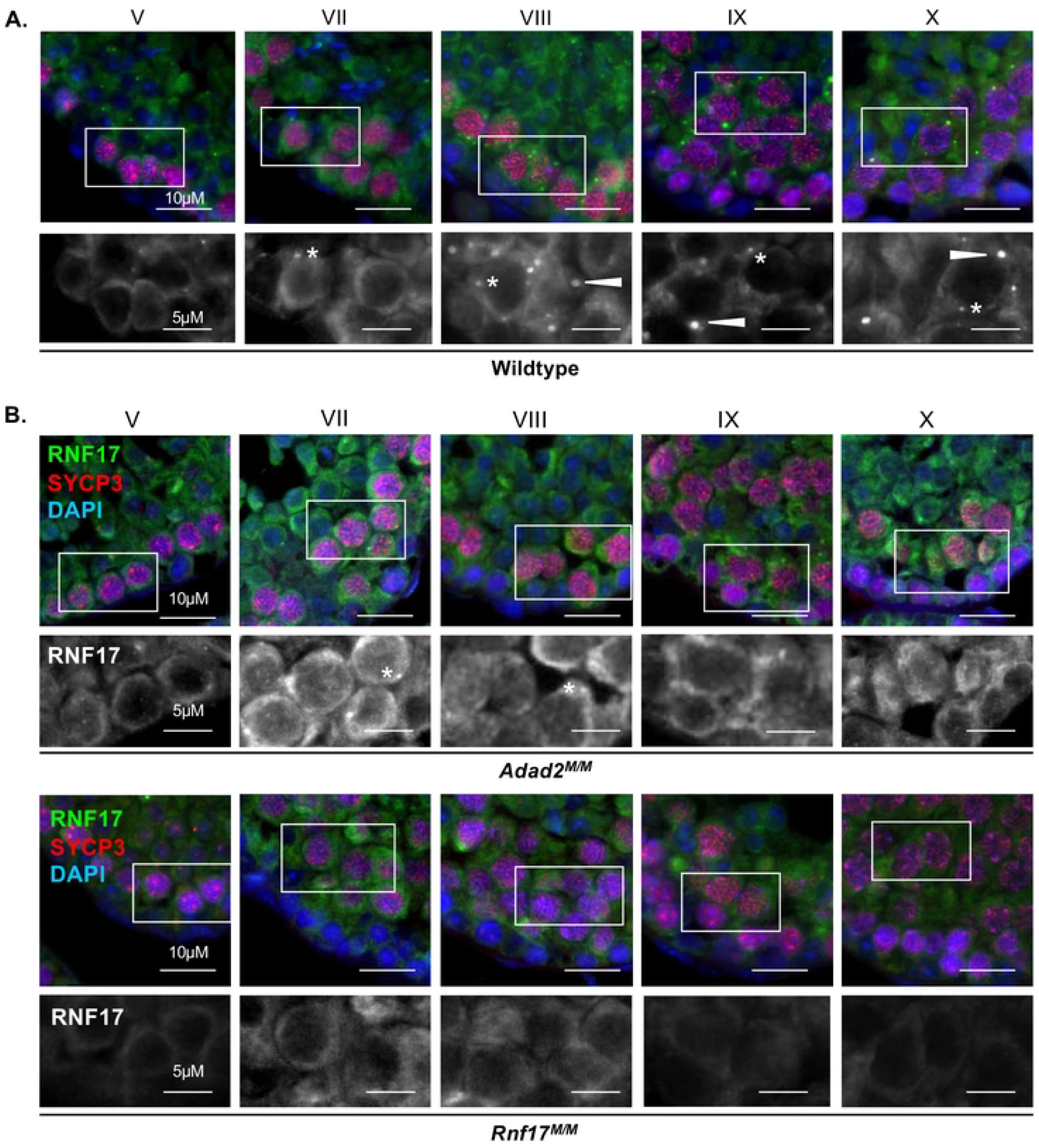
RNF17 forms distinct granules in pachytene spermatocytes and requires ADAD2 for its localization. RNF17 localization across spermatocyte development in **A**. adult wildtype testes demonstrating two phases of granule formation and two different granule types (asterisks - small granules and arrowheads – large granules) and **B**. *Adad2*^*M/M*^ and *Rnf17*^*M/M*^ mutant testes demonstrating RNF17’s reliance on ADAD2 for formation of large RNF17 granules. Roman numerals – testis tubule cross-section stage (V containing early stage pachytene spermatocytes, VII and VIII containing mid-stage pachytene spermatocytes, IX through X containing late-stage pachytene spermatocytes). Asterisks - small RNF17 granules. Green – RNF17, red – SYCP3, blue – DAPI. 400x magnification.

Given that RNF17 is still present in the absence of ADAD2 (S1A and S1B Figs), we sought to determine whether loss of ADAD2 impacted RNF17’s spermatocyte localization. Immunofluorescence of RNF17 in *Adad2^M/M^* and *Rnf17*^*M/M*^ mutant testes (Fig 3B) revealed that although cytoplasmic RNF17 along with small RNF17 granules can be detected in *Adad2^M/M^* spermatocytes, RNF17 fails to coalesce into large granules (Fig 3B), irrespective of stage. Given the overlapping timing for the ADAD2 granule and the RNF17 granule, we next examined ADAD2 granularization in the context of RNF17 loss (S3B Fig). Like RNF17, ADAD2 fails to form large granules with RNF17 loss. In addition, ADAD2 fails to form small granules in the absence of RNF17. Together, these findings suggest the ADAD2-RNF17L interaction is required for the formation of all ADAD2 granules as well as the large RNF17 granules.

### ADAD2 and RNF17 form a unique germ cell granule

ADAD2 and RNF17 share distinct phenotypic and developmental similarities. This, combined with their biochemical interaction and their reliance on one another for their granular localization, led us to wonder whether the large ADAD2 granule and the large RNF17 granule are one in the same. Immunofluorescence using fluorophore labeled anti-ADAD2 and anti-RNF17 in wildtype testes revealed near perfect colocalization between ADAD2 and RNF17 in mid-to late pachytene spermatocytes (Fig 4A). Confirmation of labeled antibody specificity was further confirmed in *Adad2* and *Rnf17* mutant testes (S4A Fig). This analysis demonstrated ADAD2 and RNF17 localize to the same granules in spermatocytes. Given this, further reference will be to the large or small ADAD2-RNF17 granules.

**Figure 4.**
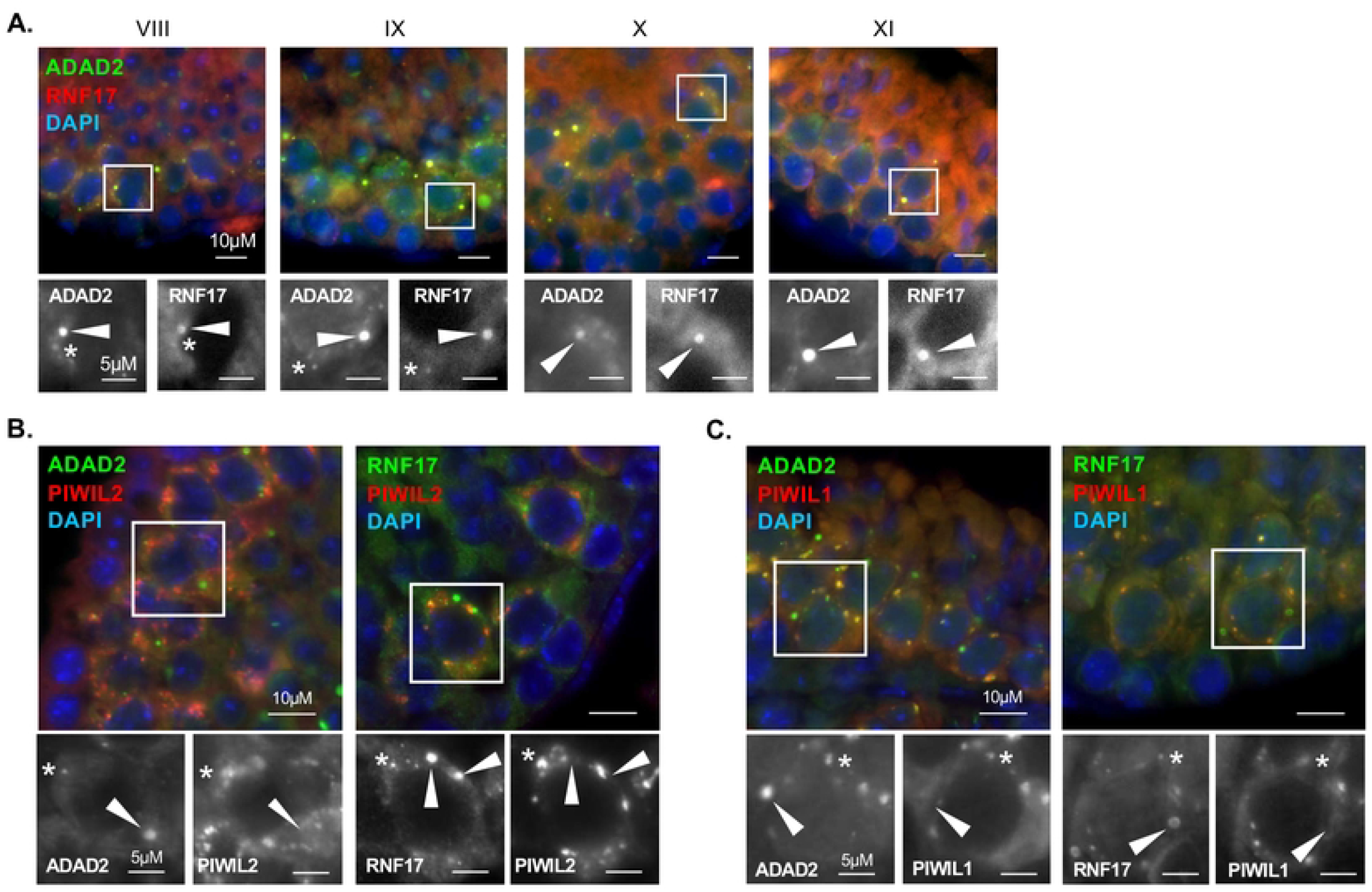
ADAD2 and RNF17 colocalize to form two distinct populations of granules. **A**. Co-immunofluorescence of ADAD2 and RNF17 in adult wildtype testes across selected pachytene spermatocyte developmental stages demonstrating colocalization. Roman numerals – testis tubule cross-section stage (VIII containing mid-stage pachytene spermatocytes, IX through XI containing late-stage pachytene spermatocytes). Asterisks - small granules and arrowheads - large granules. Red - RNF17, green - ADAD2, and blue - DAPI. Co-immunofluorescence of **B**. PIWIL2 and **C**. PIWIL1 with ADAD2 or RNF17 in adult wildtype testes demonstrating co-localization in a subset of ADAD2 or RNF17 granules. Red - PIWIL2 or PIWIL1, green - ADAD2 or RNF17, and blue – DAPI. Asterisks - small ADAD2 or RNF17 granules. Arrowheads - large ADAD2 or RNF17 granules. 630x magnification for all images.

The mammalian spermatocyte contains five distinct germ cell granules [3,24], as identified by electron microscopy (EM). These granules are crucial for proper germ cell development [5,6,31], though their exact protein composition and function are not wholly described [22]. In spite of this, several proteins are observed across four of the five granules, including DDX4 [19] and DDX25 [19,20]. The only granule known to be negative for DDX4 and DDX25 is referred to as the “cluster of 30 nm particles”, which is observed via EM from late pachytene until the end of meiosis [3,24]. Previous analysis of ADAD2 granules has demonstrated ADAD2 does not colocalize with DDX25 [23], suggesting the ADAD2 granule is unique among the protein-associated granules. However, ADAD2’s localization with DDX4 and the localization of RNF17 in relation to these markers is entirely undescribed.

To determine whether the large ADAD2-RNF17 granule represents the protein-orphan “cluster of 30 nm particles” or if ADAD2 and RNF17 instead localize with one of the better described granules, we examined ADAD2 and RNF17 granule co-localization with DDX4 along with RNF17’s co-localization with DDX25 (S4B and S4C Figs). For both ADAD2 and RNF17, no co-localization with DDX4 was observed in large or small granules. And, like ADAD2, all RNF17 granules fail to co-localize with DDX25 suggesting the ADAD2-RNF17 granule may represent the cluster of 30-nm particles. To further define the molecular composition of the ADAD2-RNF17 granule, we examined the localization of a third granule associated RBP, NANOS1 [21,24] (S4D Fig). Although NANOS1 has been reported by immuno-EM to localize to a similar set of granules as DDX4 and DDX25, NANOS1 detection by this method was shown to be extremely weak [24] and thus may not be entirely representative of true NANOS1 localization. Supporting this notion, both ADAD2 and RNF17 large and small granules were positive for NANOS1. Together, the unique molecular signature of the ADAD2-RNF17 granules defined here (negative for both DDX4 and DDX25 but positive for NANOS1) indicates they are not among the four described spermatocyte germ cell granules and thus, based on protein composition, may represent the “cluster of 30 nm particles”.

We further wondered whether the large versus small ADAD2-RNF17 granules are molecularly distinct from one another. As a measure of this, we assessed the localization of two well defined granule proteins in relation to ADAD2 and RNF17. These proteins, PIWIL1 and PIWIL2, are both associated with processing of small non-coding RNAs known as piRNAs (4,13,14) as well as localizing to granule structures in the pachytene spermatocytes [14,19]. To date, both PIWIL1 and PIWIL2 are most closely associated with large ribonuclear complexes referred to as piRNA-p-bodies [32] which can be visualized in the spermatocyte cytoplasm [33] and overlap in large part with the well-defined IMC granule [18]. Additionally, RNF17 has been implicated as a major regulator of piRNA biogenesis via interaction with PIWIL1 [34] suggesting at least a subpopulation of RNF17 granules may also contain either PIWIL1 or PIWIL2. Co-immunofluorescence was used to determine if ADAD2 and/or RNF17 localize with either PIWIL1 or PIWIL2 (Fig 4B and 4C) in the context of either the large or the small ADAD2-RNF17 granule. For both ADAD2 and RNF17, co-localization with both PIWIL1 and PIWIL2 was observed in a subset of granules. These PIWIL1 or PIWIL2 positive ADAD2 or RNF17 granules were, on average, much smaller than the large ADAD2-RNF17 granules, as such they most likely represent the pool of small ADAD2-RNF17 granules that are observed during mid-to late spermatocyte development. Given the lack of DDX4 in the small ADAD2-RNF17 granules, their colocalization with PIWIL1 and PIWIL2 suggests the small ADAD2-RNF17 granules represent a novel subtype of piRNA-containing granule, independent of the classically defined IMC. Together, these observations demonstrate the ADAD2-RNF17 granules represent two molecularly distinct populations, with the smaller pool likely representing a previously undefined subpopulation of piRNA-associated granules and the large granule representing the cluster of 30 nm particles and of unknown function.

### The large ADAD2-RNF17 granule has a unique structure with distinct domains

Initial localization studies of the large ADAD2-RNF17 granule suggested it may be roughly spherical. However, several unusual features were observed (see Fig 4B and C, individual panels for ADAD2 and RNF17) including regions of low or no signal in the center of the granule, which suggested the granule may be composed of subdomains, similar to those observed in other granule types such as stress granules [35,36]. To better define the large ADAD2-RNF17 granule structure, we first examined ADAD2 localization to determine whether it displayed variable localization within the granule (Fig 5A). This preliminary analysis revealed ADAD2 forms a distinct cup or funnel shape. The diameter of this structure can be seen as a ring, with an intense ADAD2 signal around the outer edge and a significantly weaker ADAD2 signal in the interior. These rings measure 1.036 µm ± 248 nm, (n = 24) across with the weakly positive interior region measuring 452 ± 129 nm, (n = 24). To eliminate the possibility of technical artifacts, imaging was repeated with an alternate antibody against ADAD2 (“93Term”) [23], which requires an alternate antigen retrieval method. Further, to eliminate the possibility of incomplete antibody penetration, analyses were repeated on thick wildtype slides to ensure capture of the entire structure (S. Movie 1). In all cases, ADAD2 localization appeared similar suggesting the overall shape of the granule is cup or funnel shaped with a notably sharp and flat top rim and sides. Parallel analyses were performed for RNF17 which, much like ADAD2, displayed a similar localization pattern. However, measurements of the RNF17 granule diameter demonstrated it to be somewhat larger than ADAD2 at 1.412 µm ± 591 nm, (n = 26). Likewise, the interior measurements of the RNF17 ring (558.461 ± 285.590 nm, (n = 26)) suggested the overall RNF17-dense region of the granule to be slightly larger than the ADAD2-dense region.

**Figure 5.**
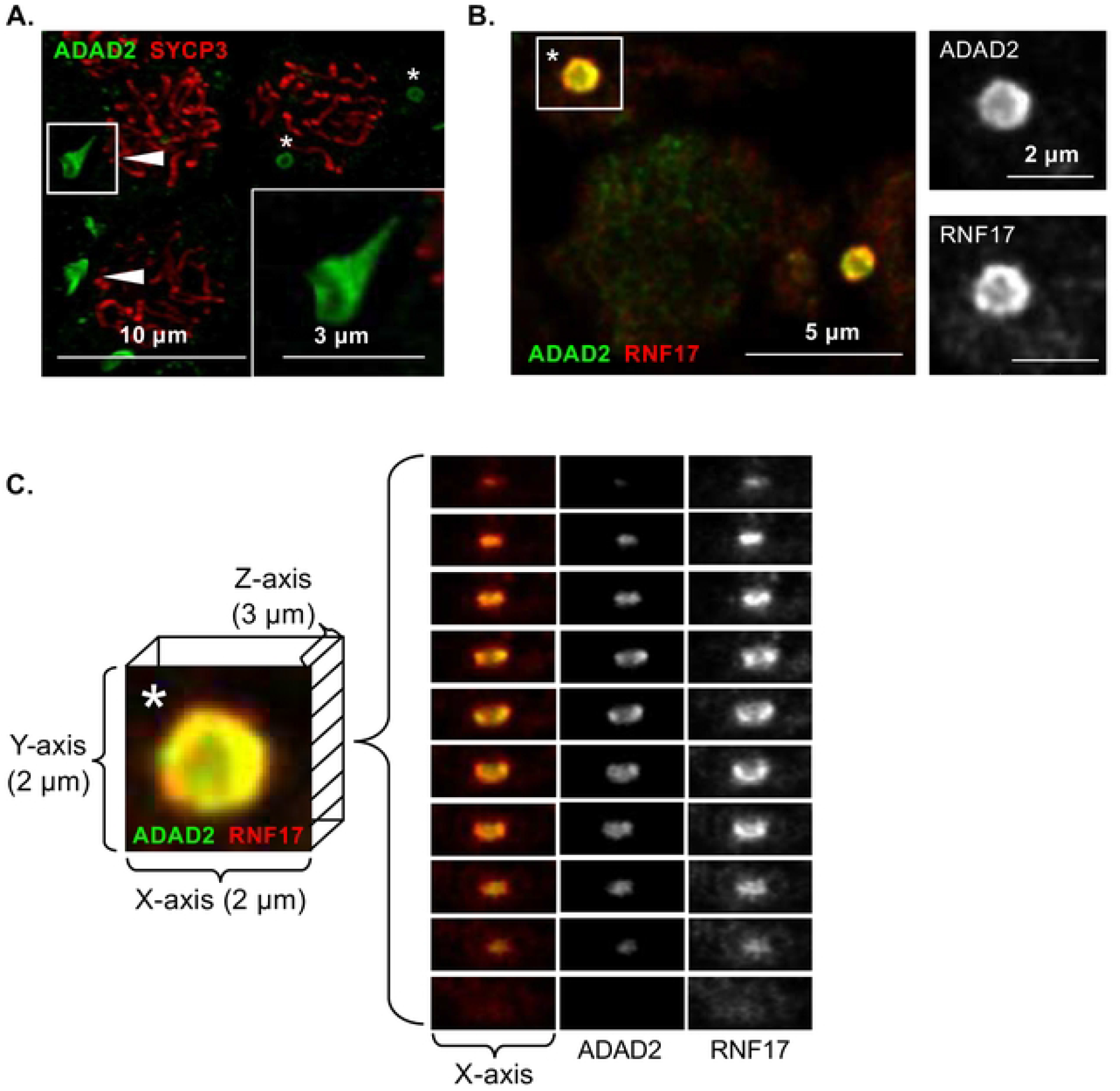
ADAD2 and RNF17 form a uniquely shaped granule. High-resolution confocal of **A**. ADAD2 in late pachytene spermatocytes counterstained with SYCP3 demonstrating a distinct cup shape structure. Red – SYCP3, green – ADAD2. 1000x magnification. **B**. Co-localization of ADAD2 and RNF17 in wildtype testis sections. Red - RNF17 and green - ADAD2. 1000x magnification. **C**. Selected ADAD2-RNF17 granule (asterisk in B) with representative XZ planes demonstrating the interior of the cup-shaped structure and the relative localization of ADAD2 and RNF17. XZ planes range from 0.13 to 0.39 µm apart. Red - RNF17 and green - ADAD2. 1000x magnification.

To shed light on whether ADAD2 and RNF17 comprise different protein domains within the granule, co-localization studies using high-resolution confocal were performed (Fig 5B, S. Movie 2). This analysis demonstrated distinct and differential localization of ADAD2 and RNF17 within the granule, with ADAD2 observed as a ring of dense aggregation around the rim and sides with much weaker signal in the interior of the cup. RNF17 was observed towards the exterior of the ADAD2 aggregation with moderate overlapping of the two domains. Further, RNF17 appeared to be even less enriched in the interior of the cup than ADAD2. This localization pattern held throughout the 3D structure of the granule (Fig 5C), with the RNF17 domain observed exterior to, but retaining the same shape of, the ADAD2 domain. Together, these findings demonstrate that the ADAD2-RNF17 granule has distinct regions, and these regions are defined by enrichment of either ADAD2 or RNF17.

### The large ADAD2-RNF17 granule is associated with the endoplasmic reticulum

No other germ cell granule is known to have the distinct shape of the large ADAD2-RNF17 granule. As germ cell RNA granules are not bound by membranes of their own [3], they are shaped by the interactions between their components and the cellular environment. The non-spherical shape of the ADAD2-RNF17 granule suggests contact with another cellular component. We therefor sought to determine if the ADAD2-RNF17 granule associates with a membrane bound organelle which may provide the surface to shape the ADAD2-RNF17 granule. Thus, we examined the co-localization of both ADAD2 and RNF17 with markers of the nuclear membrane (Lamin A/C) [37], the mitochondrial membrane (COX IV) [38], and the endoplasmic reticulum (SERCA1) [39]. Lamin was notably excluded from sites enriched for either ADAD2 and RNF17 (Fig 6A) while the co-localization of ADAD2 or RNF17 with COX IV showed a slightly more complex profile (Fig 6B). COX IV was never observed near or within the large ADAD2 or RNF17 granules. However, two populations of small ADAD2 and RNF17 granules were observed, some in close association to COX IV signal and some not. This observation further supports the notion that the small ADAD2-RNF17 granule is distinct from the large and that at least a subset of small ADAD2-RNF17 granules represent a novel, non-IMC piRNA-associated granule.

**Figure 6.**
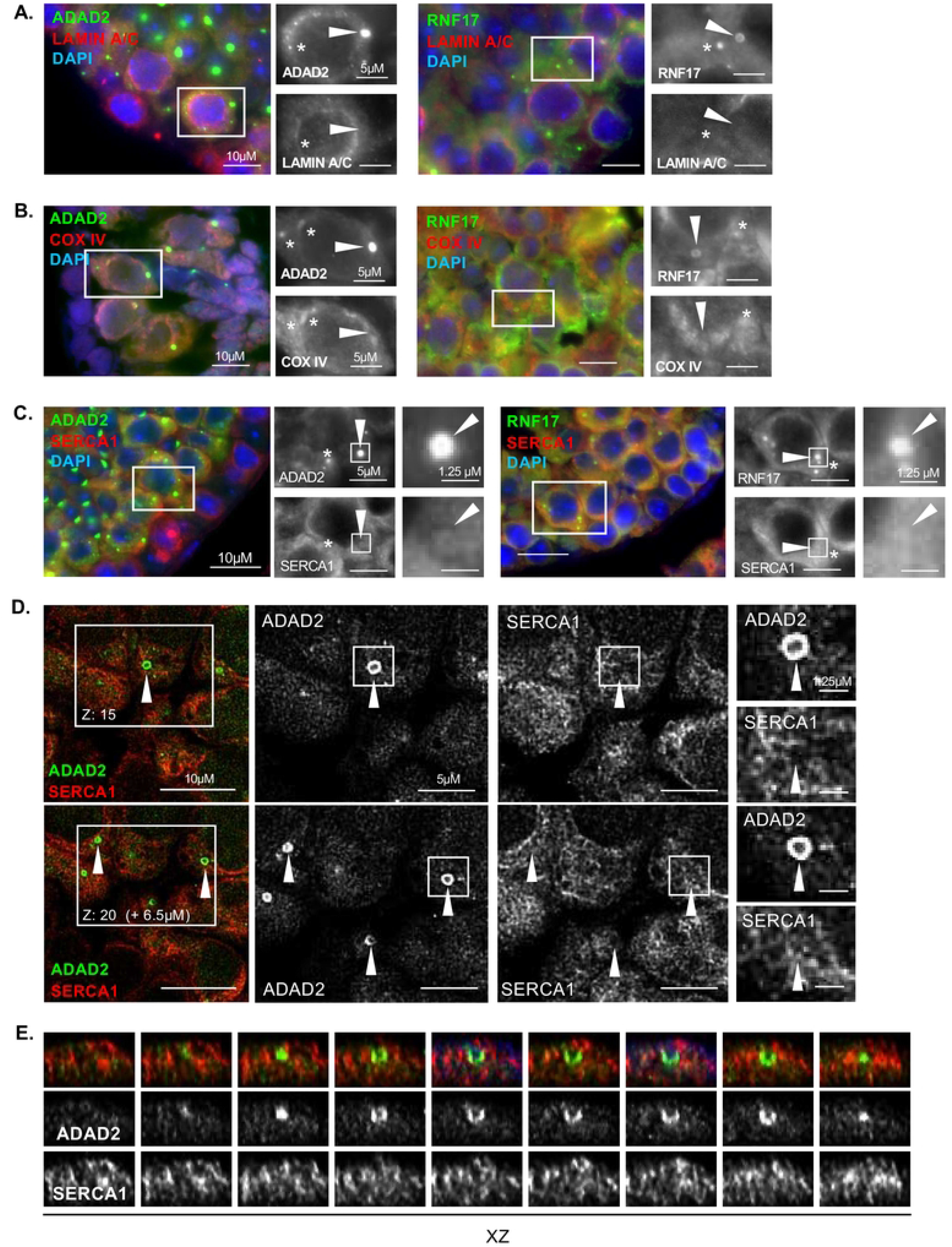
The large ADAD2-RNF17 granule associates with the endoplasmic reticulum. **A**. Co-immunofluorescence in adult wildtype testes of the nuclear membrane marker Lamin A/C or **B**. the mitochondrial marker COX IV and ADAD2 or RNF17 demonstrating neither ADAD2 nor RNF17 large granules colocalize with either the nuclear membrane or mitochondria but a subset of small granules colocalize with the mitochondria. Red – Lamin A/C or COX IV, green - ADAD2 or RNF17, and blue - DAPI. Asterisks – small ADAD2 or RNF17 grnaules and arrowheads – large ADAD2 or RNF17 granules. 630x magnification for above images. **C**. Co-immunofluorescence in adult wildtype testes of the endoplasmic reticulum marker SERCA1 and ADAD2 or RNF17 showing clustering of the SERCA1 signal at large ADAD2 or RNF17 granules. Red – SERCA1, green - ADAD2 or RNF17, and blue - DAPI. Asterisks – small ADAD2 or RNF17 grnaules and arrowheads – large ADAD2 or RNF17 granules. 400X magnification. **D**. Confocal imaging of SERCA1 and ADAD2 in adult wildtype testes showing SERCA1-enriched regions associate with the edges of the large granules. Red – SERCA1, green – ADAD2. Arrowheads – large ADAD2 granules. 1000x magnification. **E**. XZ-plane slices of the ADAD2 granule amid SERCA1-marked ER. Red – SERCA1, green – ADAD2. 1000x magnification.

In contrast to both Lamin and COX IV, SERCA1 was not excluded from large granules of either ADAD2 or RNF17 and occasional regions of SERCA1 and ADAD2 or RNF17 co-enrichment detected (Fig 6C) suggesting potential association of the ER with the large ADAD2-RNF17 granule. To better define the spatial association of the endoplasmic reticulum with the ADAD2-RNF17 granule, we examined the co-localization of ADAD2 and SERCA1 using confocal microscopy. These analyses clearly identified the distinct localization of the ADAD2-RNF17 granule along with defining the tubular structure indicative of the ER [40] (Fig 6D). Unlike our previous analyses, this analysis revealed a distinct void in the ER where the ADAD2-RNF17 granule resided. However, these voids were surrounded with SERCA1, likely driving the apparent overlap in localization observed with traditional IF imaging. Examination of the XZ plane (Fig 6E) demonstrated association of SERCA1 along the exterior of the ADAD2-RNF17 granule with minimal co-localization within it. Together, these results suggest the unique shape of the ADAD2-RNF17 granule is due to its association with the endoplasmic reticulum, though the driving factors behind this association remain unclear.

### Loss of both ADAD2 and RNF17 phenocopies the *Adad2* and *Rnf17* phenotype

To assess whether the loss of both ADAD2 and RNF17 had a more profound impact on germ cell development than singular loss, *Adad2*^*M/M*^*:Rnf17*^*M/M*^ mice were generated. Dual loss of both ADAD2 and RNF17 was confirmed via Western blot (S5A Fig). Though female *Adad2*^*M/M*^*:Rnf17*^*M/M*^ (double mutant) mice were fertile, males were completely infertile and demonstrated post-meiotic germ cell loss, with variable severity between tubules (S5B Fig). When round spermatids were assessed in double mutant males, a significant decrease was observed compared to wildtype (S5C Fig), similar to either ADAD2 or RNF17 loss. Further, examination of the heterochromatin landscape in double mutant round spermatids demonstrated heterochromatin abnormalities consistent with those observed in both *Adad2*^*M/M*^ and *Rnf17*^M*/M*^ single mutants (Fig 7A). Quantitation of these abnormalities ultimately revealed that the double mutant males exhibited similar frequencies of abnormal H3K9me3 foci as observed in both single mutants (Fig 7B). As the phenotype of the double mutants mimics that of either single mutant, these results suggest that the formation of the ADAD2-RNF17 granules, or the proteins’ interaction, is more crucial to successful germ cell development than the presence of the proteins themselves.

**Figure 7.**
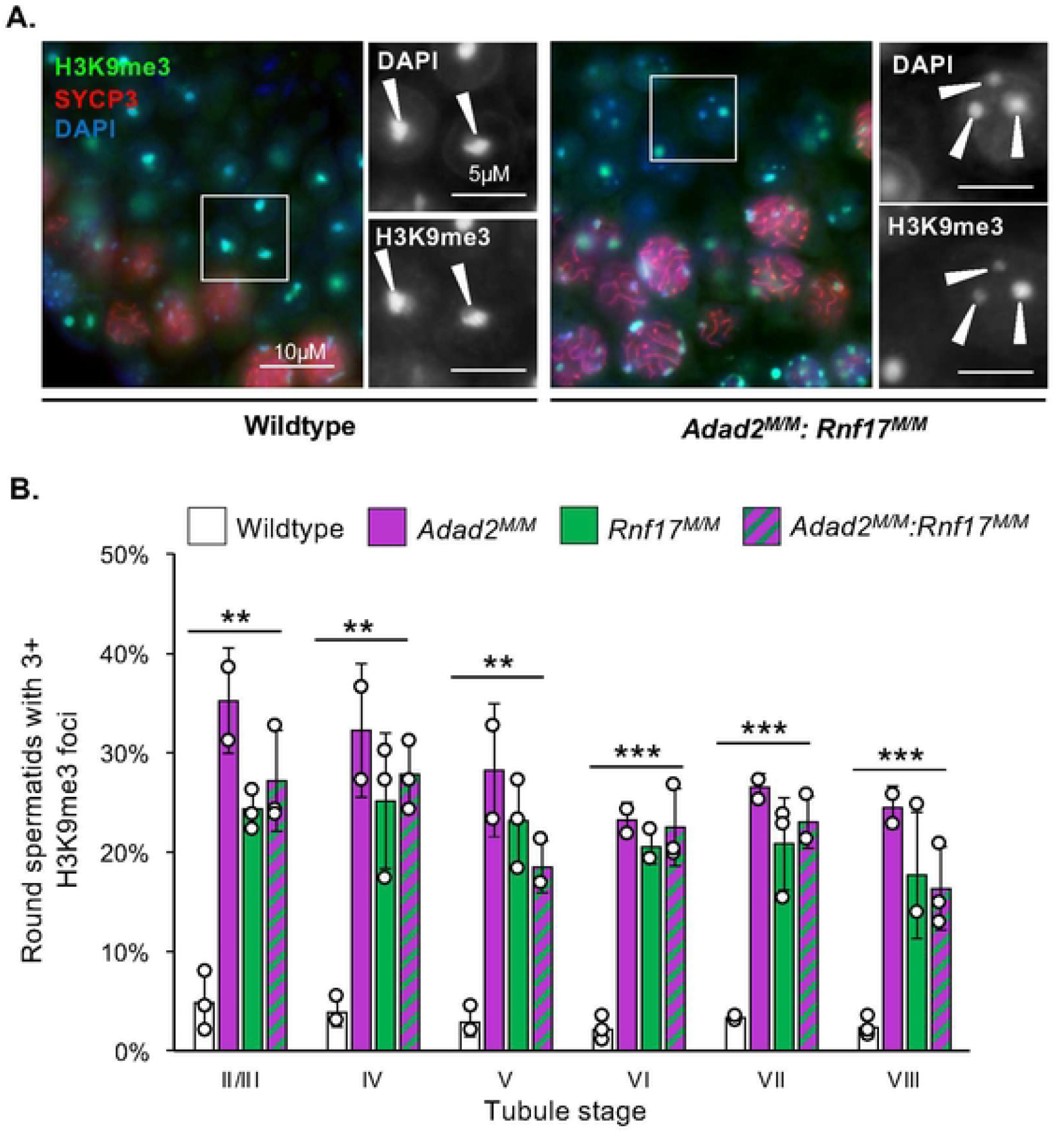
*Adad2:Rnf17* double mutants have a very similar chromocenter phenotype to that observed in single mutants. **A**. Immunofluorescence of the heterochromatin mark H3K9me3 along with DAPI staining in adult wildtype and *Adad2*^*M/M*^: *Rnf17*^*M/M*^ testes. Red - SYCP3, green - H3K9me3, and blue - DAPI. Arrowheads – DAPI or H3K9me3 foci in round spermatids. 400x magnification. **B**. Quantification of round spermatids with 3 or more chromocenter-like structures as marked by H3K9me3 staining per developmental stage in adult wildtype, *Adad2*^*M/M*^: *Rnf17*^*M/M*^, *Rnf17*^*M/M*^ (n = 3), and *Adad2*^*M/M*^ (n = 2) testes demonstrating a significant increase in *Adad2*^*M/M*^: *Rnf17*^*M/M*^ round spermatids relative to wildtype. Data are mean ± s.d. Significance was calculated using an unpaired, one-tailed Student’s t-test between wildtype and *Adad2*^*M/M*^: *Rnf17*^*M/M*^ (\**P* < 0.05, \*\**P* < 0.005, \*\*\**P* < 0.0005).

## Discussion

In mammalian meiotic male germ cells, germ cell granules are important sites of RNA metabolism required for their normal differentiation. They contain at least five morphologically distinct granule types and loss of many granule components leads to male infertility. In spite of this, there is limited knowledge regarding the protein composition of, function of, and relationship between these granules. To that end, this work aimed to characterize a recently identified granule that contains the RNA binding protein ADAD2, which is required for male fertility [23]. This work leveraged protein interaction studies to identify a second RNA binding protein also required for male fertility, RNF17 [8], as an ADAD2 interacting partner. Genetic knockout models combined with localization studies demonstrated ADAD2 and RNF17 are co-dependent on one another to form at least two distinct populations of ADAD2-RNF17 granules. Protein composition studies of these ADAD2-RNF17 granules further showed molecularly distinct subpopulations not related to the best characterized granule, the IMC (Fig 8). Lastly, dual loss of both ADAD2 and RNF17 via genetic ablation suggested loss of the granules themselves, as opposed to the individual proteins, is the primary driver of the *Adad2* and *Rnf17* mutant phenotypes. Together, these studies have genetically defined the importance of multiple novel germ cell granules in male fertility and have additionally revealed new aspects of germ cell granule biology that establish future directions and approaches for the study of other germ cell granules.

**Figure 8.**
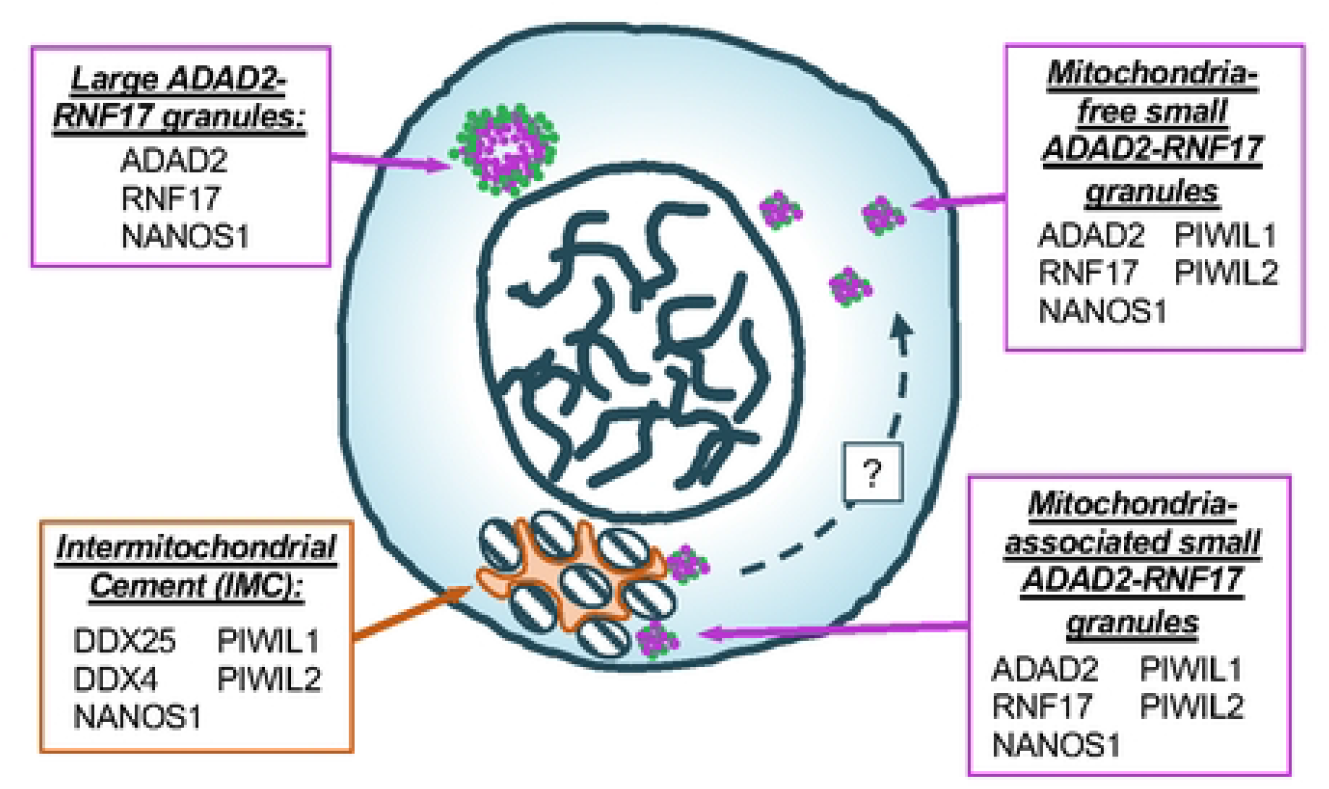
Model of the relationship between ADAD2-RNF17 granules and the IMC. Three populations of ADAD2-RNF17 granules are observed in meiotic spermatocytes. Of these, one is associated with the mitochondria but does not share many protein components of the IMC. The remaining two are molecularly distinct from one another. There is a potential relationship between IMC-associated and non-IMC associated ADAD2-RNF17 granules, though that relationship is currently unclear.

Most genetic studies of male germ cell granules to date have utilized single gene knockout models [11,31,41–43]. Although powerful for defining the function of the targeted protein, these approaches fail to inform on the function of the granules themselves as they also impact any non-granule functions of the chosen target [22]. Supporting this notion, single gene knockouts of granule-associated proteins often lead to phenotypes prior to formation of the granule [13,31]. Single loss of ADAD2 and RNF17 avoids these complications as both result in phenotypes after granule formation. Further, their reliance on one another for granule formation means reciprocal studies in the two models provide an opportunity to study the granule-specific function for each protein while still leaving cytoplasmic pools available to carry out any necessary functions. Given both models demonstrate a striking similar phenotype, we surmised the granule itself was important for ADAD2- and/or RNF17-driven physiology. Supporting this notion, compound mutation of both *Adad2* and *Rnf17* gave rise to a quantitatively similar phenotype as single mutation. These findings lead to a paradigm for testing the function of germ cell granules themselves that requires only identification of granule-associated binding partners necessary for granularization of individual proteins. One especially promising class of RNA binding proteins that meet these criteria are the Tudor-domain containing protein family (TDRDs) [13,44], of which RNF17 is a member [8]. Several TDRDs have already been shown to tether granule components to their respective granules [44], providing a proof of concept for this approach. Importantly, as Tudor-domains have well-defined protein binding preferences based on post-translational modifications [44–46], their study may also provide valuable insight into the direct molecular mechanisms driving germ cell granule formation, an area for which there is very limited knowledge.

The demonstration that the two pools of ADAD2-RNF17 granules are molecularly distinct from one another has several important implications, not only for understanding their biology, but the biology of other germ cell granules as well. First, based on the protein composition of the ADAD2-RNF17 granules described herein, we conclude both granules likely represent unstudied granule populations in the germ cell. Concerning the large ADAD2-RNF17 granule, its size and apparent density makes it likely to be detected by EM [3]. Further, lack of co-localization with the granule protein DDX4 strongly suggests it is the protein-orphan granule “cluster of 30 nm particles”, which is the only granule known to be negative for DDX4 [24]. As such, this report represents the first identification of proteins associated with this exclusively EM-defined structure. Further, localization of NANOS1 to the structure provides the first hint towards a potential function. The *Drosophila melanogaster* homolog of NANOS1 is known to facilitate localized translation regulation in the embryo [47] while in *Xenopus*, Nanos1 facilitates translational repression in the germline [48]. Together, the emerging model is that NANOS1 may play important roles in regulating translation. Whether it does so in the context of the ADAD2-RNF17 granule remains an open and exciting avenue of study.

That NANOS1 appears to be shared between the large ADAD2-RNF17 granule and the other EM-defined granules is also the first biological connection between the orphan granule and the broader granule population. For the first time, this suggests that all the EM-defined granules may share one or more components and calls into question the commonly held belief that germ cell granules represent distinct pools within the germ cell cytoplasm. This notion is supported by co-localization studies of the small ADAD2-RNF17 granule. As all small ADAD2-RNF17 granules contain the piRNA processing proteins PIWIL1 and PIWIL2, they are likely associated with either piRNA processing or piRNA action. To date, the IMC is the only known site for piRNA processing in meiotic male germ cells [14]. However, a subset of small ADAD2-RNF17 granules is separate from mitochondria, demonstrating them to be independent of the IMC. These observations make it tempting to postulate the small ADAD2-RNF17 granules represent sites of different piRNA processing or action steps. Together, the identification of distinct ADAD2-RNF17 granule pools brings into question our understanding of other granules and suggests that rather than distinct units they instead may represent an interconnected network of RNA biogenesis or regulation sites. Several previous observations support this hypothesis. First, as discussed above and recently reviewed [22], there are multiple proteins observed across many granules. Second, multiple granules have already been shown, by both EM and immunodetection, to actively share components [3,19,24,49]. Although this hypothesis will require continued rigorous testing, it represents an exciting future avenue of germ cell granule research.

The last findings of importance to the broader germ cell granule field herein are those focused on the structure of the ADAD2-RNF17 granule. First, localization of ADAD2 within the large granule identified distinct regions of high and low density, in particular along the rim of the structure and relative to the interior. This is reminiscent of what is observed in stress granules, especially for the stress granule protein G3BP1 [50–52]. Stress granules are well-described, RNA-rich, non-membrane bound structures that form under specific cellular stresses [53]. As such, they represent an excellent model system to better understand granule formation. Recent work has demonstrated that nucleation of G3BP1 drives stress granule formation, which is ultimately the result of liquid-liquid phase separation [54]. It is unknown whether this is a primary driver of ADAD2-RNF17 granule formation or the formation of other germ cell granules. However, it is known that DDX4 undergoes phase separation in vitro [55] and multiple groups have proposed phase separation to be the primary driver of granule formation in vivo [56–59].

Although it seems likely the ADAD2-RNF17 granule is formed, in part, by phase separation other observations suggest additional levels of regulation. In contrast to stress granules, which are spherical or oblong in nature and are not known to interact with any specific membrane-bound organelles [56], the ADAD2-RNF17 granule displays multiple facets that appear flat. This suggests some sort of physical or mechanical constraint on granule formation, perhaps similar to the constraints mitochondrial tethering puts on IMC components [44,60]. Supporting this, the ADAD2-RNF17 granule appears to be in close contact with the ER membrane, which may be providing the necessary mechanical force to generate the flat surfaces observed in the ADAD2-RNF17 granule. Whether ADAD2 and/or RNF17 are directly or indirectly tethered to the ER remains a question for future work.

In total, this work has identified several protein components of an entirely undescribed germ cell granule population as well as defined the granules themselves as necessary drivers of male germ cell differentiation. Importantly, this work lays the foundation to address multiple outstanding questions regarding the ADAD2-RNF17 granules. These include the exact molecular function of the granule subpopulations, the purpose of the protein subdomains within the large granule structure, and whether other associated proteins drive one or both of these. Study of the ADAD2-RNF17 granules has already provided novel insight into the biology of other germ cell granules, As such, addressing these future questions should not only inform on ADAD2 and RNF17 biology but on other germ cell granules as fundamental drivers of male germ cell differentiation.

## MATERIALS AND METHODS

### Animal care and model generation

All animal use protocols were approved by the Rutgers University animal care and use committees. Mouse procedures were conducted according to relevant national and international guidelines as outlined in the Guide for the Care and Use of Laboratory Animals and provisions set forth by the Animal Welfare Act. Adherence to these guidelines was overseen on the institutional level by the Rutgers Institutional Animal Care and Use Committee and all animal procedures were approved under Rutgers IACUC ID TR202000034. Generation of *Adad2*^*M/M*^ mice was as described by Snyder et al. [23] and *Rnf17*^*M/M*^ mice were as described by Pan and colleagues [8]. *Adad2:Rn17* mice were generated by breeding *Adad2*^*M/M*^ females to *Rnf17*^*+/M*^ males as well as *Rnf17*^*M/M*^ females to *Adad2*^*+/M*^ males to generate *Adad2*^*+/M*^: *Rnf17*^*+/M*^ offspring. These were intercrossed to generate *Adad2*^*M/M*^: *Rnf17*^*M/M*^, *Adad2*^*+/+*^: *Rnf17*^*M/M*^, *Adad2*^*M/M*^: *Rnf17*^*+/+*^ and *Adad2*^*+/+*^: *Rnf17*^*+/+*^ experimental animals. All mice were housed in a sterile, climate-controlled facility on a 12 h light cycle. Mice were fed LabDiet 5058 irradiated rodent chow and had access to food and water ad libitum.

### Immunoprecipitation

Testes were collected from 42 dpp *Adad2*^*M/M*^, *Rnf17*^*M/M*^, and wild-type mice (n = 3) and flash frozen. Tissue was ground in liquid nitrogen and total protein extracted in RIPA buffer (50mM Tris-HCl (pH 8), 150 mM NaCl, 1% (v/v) NP-40, 0.5% (w/v) sodium deoxycholate, 0.1% (w/v) SDS) with protease inhibitors (Thermo Scientific, 1 tablet per 10 mL) at a ratio of 1 ml buffer to 100 mg tissue. Tissue was precleared with Protein A beads (Thermo-fisher) equilibrated in RIPA (10 µL beads per 100 mg tissue) for one hour at 4°C. For every 1 mL of lysate, 4 µL of either ADAD2 antibody (94 AP [23]) or 2.5 µg RNF17 antibody (Proteintech) was added. Samples were incubated with antibody overnight at 4°C with constant rotation. Subsequently, beads were added and left to bind for 2 hours at 4°C with rotation followed by bead washing with 0.25X TBS (5 mM Tris Base, 37.5 mM NaCl). For mass spectrometry, protein was extracted from beads with 30 µL glycine elution buffer (0.2 M glycine, pH 2.6) per 1 ml of lysate followed by neutralizing with Tris pH 8.0. Glycine elution was confirmed via SYPRO gel staining (as per manufacturer’s protocol, see below) and western blotting (S6 Fig). Analysis of IP protein by mass spectrometry was performed as below. For western blot confirmation of identified interactions, IP beads were boiled for 5 minutes at 95°C in SB loading dye (30 µL dye per 1 mL). Eluent was then isolated from the beads and prepared for western blotting along with controls (input, flow-through, and first wash).

### Mass spectrometry

Protein eluents for IP-MS were digested with trypsin after resuspension in 20 mM Tris-HCl pH 8.0. Samples were reduced with DTT and alkylated with iodoacetamide prior to addition of trypsin for overnight digestion at 37 °C followed by quenching with formic acid and salt removal by p10 ZipTip. Samples were analyzed at Northwestern University Proteomics on a ThermoFisher Orbitrap. Data was then analyzed by the Mass Spectrometry and Protein Chemistry Service at The Jackson Laboratory using MASCOT against the SwissProt 2015_08 Mus musculus database. Fixed modifications were set for carbamidomethyl and variable modifications for acetyl/phosphorylation. Peptide mass tolerances were set to 25 ppm and 0.2 Da for intact and fragments, respectively. Proteins of interest were identified by comparison of wildtype and mutant IP samples.

### Protein isolation and Western blotting

Testes were collected from adult (60-70 dpp) and 21 dpp *Adad2*^*M/M*^, *Rnf17*^*M/M*^, and wildtype male mice and flash frozen. Tissue was ground in liquid nitrogen and total protein was extracted by RIPA buffer with protease inhibitors at a ratio of 1 ml buffer to 100 mg tissue. Protein concentration was determined via the DC protein assay (BioRad) as per manufacturer’s instructions. Samples were diluted in RIPA buffer, and 1M dithiothreitol and loading dye (100 mM Tris-HCl (pH 6.8), 4% (w/v) SDS, 0.05% (w/v) bromophenol blue, 20% (w/v) glycerol) was added in a ratio of 2:1:2. Samples were boiled at 95°C for 5 minutes prior to loading.

For Western blotting of total protein, 20 µg protein per sample was electrophoresed on 10% acrylamide gels for ADAD2 detection and 50 µg protein per sample was electrophoresed on 6% acrylamide gels for detection with RNF17. For Western blotting of IP panels, 20 µL each of input, flow-through, and wash along with 20 µL of IP diluted 1:10 was loaded per well.

Following wet transfer of proteins to a PVDF membrane (BioRad), membranes were blocked and incubated overnight with primary antibody at 4°C (anti-ADAD2 1:1000; anti-RNF17 (Proteintech) 1:1000). Blots were incubated in secondary antibody (Goat-Anti-Rabbit HRP, (Biorad) 1:2000) for one hour at room temperature. HRP detection was performed using SuperSignal West Pico PLUS Chemiluminescent Substrate (Thermo Scientific) and visualized using an Azure Biosystems C600 imager. Equal loading and transfer was confirmed after probing via SYPRO-Ruby staining (see below and S7 Fig). For a detailed breakdown of antibody concentrations and conditions, see Table S1.

### Western blot membrane and gel staining

Following visualization of blot, membranes were stained with SYPRO-Ruby Protein Gel Stain (Lonza Rockland) to confirm equal loading. Membranes were stained following manufacturer’s instructions (Molecular Probes) and visualized using an Azure Biosystems C600 imager set to UV 302. For IP gel staining, gels were processed as per manufacturer’s instructions after electrophoresis and imaged with the same method as for membranes.

### Hematoxylin and Eosin (H & E) staining

Testes from 60-70 dpp *Adad2*^*+/M*^:*Rnf17*^*+/+*^ and *Adad2*^*M/M*^:*Rnf17*^*M/M*^ males were collected and fixed overnight in Bouins Solution (Sigma Aldrich). Tissue was rinsed in deionized water, dehydrated in increasing concentrations of ethanol, embedded in paraffin wax and cut into 4 µm sections. Slides were deparaffinized in xylenes and rehydrated in decreasing concentrations of Ethanol before staining with Harris Hematoxylin (Sigma Aldrich). Slides were rinsed in water and partially dehydrated before staining with Eosin Y (Sigma Aldrich). slides were then dehydrated and mounted with Permount mounting medium (Sigma Aldrich). Samples were visualized on a custom-built microscope (Zeiss) with fluorescent and brightfield capabilities.

### Round spermatid quantification

Testes from adult wild-type, *Adad2*^*M/M*^, *Rnf17*^*M/M*^, *Adad2*^*M/M*^:*Rnf17*^*+/+*^, *Adad2*^*+/+*^:*Rnf17*^*M/M*^ and *Adad2*^*M/M*^:*Rnf17*^*M/M*^ (double mutant) testes were collected and fixed overnight in 4% PFA. Tissue was rinsed in PBS, dehydrated in increasing concentrations of ethanol, embedded in paraffin wax and cut into 4 µm sections. Slides were deparaffinized in xylenes and rehydrated in decreasing concentrations of Ethanol before staining with DAPI Fluoromount-G (Southern Biotech).

Histological parameters previously described [61] and staging criteria (below) were used to quantify the number of round spermatids per tubule and number of round spermatid-containing tubules per sample. Round spermatid morphology was quantified in DAPI-stained samples. For total round spermatids, ten tubules per quantified stage were assessed per biological replicate (n = 3). For chromocenter analyses, spermatids were binned by number of intense H3K9me3-staining structures (1, 2, or 3+). Two hundred round spermatids per stage per biological replicate (n = 3) were counted. For both counts, totals and averages (means) for each genotype were calculated, as well as s.d. An unpaired, one-tailed Student’s t-test was used to identify significant differences by genotype.

### Immunofluorescence

Testes were dissected from adult mice and fixed overnight in 4% (w/v) PFA in PBS. Tissue was rinsed in PBS and dehydrated in increasing concentrations of ethanol before embedding in paraffin wax. Applications used 4 µm sections. Antigen retrieval was performed by boiling slides in Tris-EDTA pH 9.0 (10 mM Tris-HCl, 1 mM EDTA, and 0.05% Tween) on low power for 30 min or 15.9 mM Citrate pH 5.95, on high power for 2 minutes, medium power for 7 minutes and 20 minutes at room temperature (93-Term only [23]). Slides were mounted using DAPI Fluoromount-G and stored at 4°C with light protection.

Slides were visualized on a custom-built microscope (Zeiss) with fluorescent and bright-field capabilities. Each channel was imaged individually through MetaMorph imaging software (Molecular Devices) and color-combined using the built-in color combine tool. Provided images are representative of three or more biological samples. Signal intensity was matched across slides by matching background (interstitial) signal intensity. Developmental stages were determined according to the parameters set forth by Russel et al. [61], facilitated by SYCP3 co-staining where possible as outlined below. All quantification was carried out via direct visualization.

### Antibody labeling

For colocalization studies, ADAD2, RNF17, and PIWIL1 were fluorescently labeled using Zenon™ Rabbit IgG Labeling Kits (Alexafluor 488 and Alexafluor 594, Thermofisher Scientific) as per manufacturer’s instruction.

### Confocal visualization

Slides prepared for confocal visualization were processed as standard IFs up until mounting, with the same antigen retrievals and primary and secondary antibody concentrations (See above and Table S1). Applications used 4 µm sections (or 8 µm only as specified). After incubation with secondary antibody, slides were counterstained with DAPI (Sigma-Aldrich dissolved in deionized water (20 µg/µL) and applied to tissue sections and incubated in a light-protected humid chamber for 20 minutes at room temperature. Light protected slides were then rinsed in running deionized water for 20 minutes and mounted with ProLong Glass Antifade mountant (Thermo Scientific) per manufacturer’s instructions.

Slides were imaged on a Leica TCS SP8 tauSTED 3X with Lightning capabilities using the 100x objective (HC PL APO CS2 100x/1.40 OIL). Images were taken of tubules at stages IX-XI [61]. Acquisition format was 1024×1024, speed 400hz, and a pinhole of 0.5. Line average was 4 for all channels. Z-stack step size was 0.13 µm. Z-stacks were captured with Lightning using Leica Application Suite X (LAS-X, Leica) and granule measurements were taken using the inbuilt quantification functions in LAS-X. Images and movies were color-combined and re-sliced using FIJI (Image J) [62].

### Tubule staging criteria

Stages of seminiferous tubule sections were determined according to the definitions outlined previously [61] along with a combination of SYCP3 and DAPI staining. These definitions are further outlined in S8 Fig. As both *Adad2*^*M/M*^ and *Rnf17^M/M^* males do not complete spermatogenesis, staging was reliant on cell types present prior to ADAD2 expression, primarily preleptotene, leptotene, zygotene, and early pachytene spermatocytes.

## ACKNOWLEDGEMENTS

The authors would like to thank current and previous members of the Snyder laboratory including Kelly Seltzer, Christopher Eddy, Gabriella Acoury, Gabrielle Vittor, and Megan Forrest for their support with animal husbandry, molecular analyses, and critical evaluation throughout the project. We would additionally like to thank the Human Genetics Institute of New Jersey Imaging Core for their support in confocal imaging and Dr. Jessica Shivas and Daniel Jung for their assistance with confocal data acquisition and analyses. The authors would especially like to thank our funding sources: the Eunice Kennedy Shriver National Institute of Child Health and Human Development (NIH-NICHD F32 HD072628, K99/R00 HD083521, and R01 HD107066 to ES) and Rutgers University (to ES).

## SUPPLEMENTAL FIGURE LEGENDS

**Supp. Figure 1. Mutation of *Adad2* or *Rnf17* leads to minimal abundances changes of the other**. Western blot of ADAD2 or RNF17 in **A**. 42 dpp wildtype, *Adad2*^*M/M*^, and *Rnf17*^*M/M*^ whole testis (n = 3) demonstrating complete ADAD2 or RNF17 ablation in the respective genetic model and **B**. 21 dpp wildtype, *Adad2*^*M/M*^, and *Rnf17*^*M/M*^ whole testis (n = 3). Asterisks - RNF17 protein isoforms, large – RNF17L and small – RNF17S. For loading controls, see S7 Fig.

**Supp. Figure 2. ADAD2 and RNF17 share a similar developmental profile. A**. Western blot of ADAD2 and RNF17 in wildtype whole testis protein across neonatal and juvenile developmental time points demonstrating the similar developmental profile of ADAD2 and RNF17L. Asterisks - RNF17 protein isoforms, large – RNF17L and small – RNF17S. For loading controls, see S7 Fig. **B**. Immunofluorescence of ADAD2 or RNF17 in wildtype testis across juvenile development. Images represent most mature seminiferous tubule sections at each age and demonstrate small and large ADAD2 and RNF17 granules forms by 15 dpp. Asterisks – small ADAD2 or RNF17 granules. Arrowheads – large ADAD2 or RNF17 granules. 400x magnification.

**Supp. Figure 3. ADAD2 granular localization in pachytene spermatocytes requires RNF17. A**. Immunofluorescence of ADAD2 across pachytene spermatocyte development in adult wildtype testes demonstrating small and large granule formation in mid-stage pachytene spermatocytes. **B**. ADAD2 immunofluorescence in *Adad2*^*M/M*^ and *Rnf17*^*M/M*^ developing pachytene spermatocytes. Note the non-specific ADAD2 signal observed in *Adad2* mutant round spermatids. Roman numerals – testis tubule cross-section stage (V containing early-stage pachytene spermatocytes, VII and VIII containing mid-stage pachytene spermatocytes, IX and X containing late-stage pachytene spermatocytes). Asterisks - small granules and arrowheads - large granules. Red - SYCP3, green - ADAD2, and blue - DAPI. 400x magnification.

**Supp. Figure 4. Large and small ADAD2-RNF17 granules are molecularly distinct from one another and large ADAD2-RNF17 granules are unique from other defined granules. A**. Co-immunofluorescence of ADAD2 and RNF17 using fluorophore labeled anti-ADAD2 and anti-RNF17 in *Adad2*^*M/M*^ and *Rnf17*^*M/M*^ adult testes demonstrating weak, cytoplasmically diffuse non-specific signal. Red – RNF17, green – ADAD2, and blue – DAPI. **B**. Immunofluorescence of DDX4 and ADAD2 or RNF17 in adult wildtype testes demonstrating neither ADAD2 nor RNF17 colocalize with DDX4 in either granule type. Red - DDX4, green - ADAD2 or RNF17, and blue - DAPI. **C**. Immunofluorescence against DDX25 and RNF17 demonstrates that RNF17 does not colocalize with DDX25 in either large or small granules. Red - DDX25, green - RNF17, and blue - DAPI. **D**. Immunofluorescence of NANOS1 and ADAD2 or RNF17 demonstrating NANOS1 co-localization in both large and small ADAD2 or RNF17 granules. Red - NANOS1, green - ADAD2 or RNF17, and blue - DAPI. Asterisks - small granules and arrowheads - large granules. All images 630x magnification.

**Supp. Figure 5. *Adad2:Rnf17* double mutants display post-meiotic germ cell loss. A**. Western blot of 21 dpp wildtype and *Adad2*^*M/M*^*:Rnf17*^*M/M*^ whole testis lysate (n = 3) probed for ADAD2 and RNF17 confirming ablation of both proteins. **B**. Adult *Adad2*^*+/M*^: *Rnf17*^*+/+*^ and *Adad2*^*M/M*^: *Rnf17*^*M/M*^ testis tubule cross-sections stained with H&E demonstrating the range of tubule defects in double mutant testes, including significant post-meiotic germ cell loss. **C**. Average number of round spermatids per tubule per developmental stage in adult testes from wildtype and *Adad2*^*M/M*^: *Rnf17*^*M/M*^ animals (n = 3). Data are mean ± s.d. Significance was calculated using an unpaired, one-tailed Student’s t-test (\**P* < 0.05, \*\**P* < 0.005, \*\*\**P* < 0.0005).

**Supp. Figure 6. Gel and Western blot confirmation of IP efficiency in representative wildtype and *Adad2***^***M/M***^ **immunoprecipitation (IP) samples. A**. SYPRO-Ruby stained SDS-PAGE gel. **B**. Western blot against ADAD2 in 42 dpp wildtype and *Adad2*^*M/M*^ IPs.

**Supp. Figure 7. Western blot loading controls**. SYPRO-Ruby stained membranes for blots shown in **A**. S1 Fig, **B**. S2 Fig, and **C**. S5 Fig showing equal loading across lanes.

**Supp. Figure 8. Rubric for SYCP3-immunofluorescence based staging in adult testis**. Stage indicated by Roman numerals. I-II/III: SYCP3 stains small pachytene spermatocytes (*) with threads and multiple large blobs of signal and strongly stains the chromocenter of round spermatids. IV: SYCP3 stains small pachytene spermatocytes similar to II/III and the chromocenter of round spermatids, but weakly. V: SYCP3 stains larger pachytene spermatocytes with distinct threads and a few large blobs of signal. VI: SYCP3 weakly stains the B-spermatogonia (*) and large pachytene spermatocytes with distinct threads. VII: SYCP3 stains preleptotene spermatocyte nucleoplasm (*) and large pachytene spermatocytes with distinct threads. VIII: SYCP3 strongly stains preleptotene spermatocyte nucleoplasm along with intensely stained patches (*) and larger pachytene spermatocytes with distinct threads that are further apart than in VII. IX: SYCP3 strongly and unevenly stains leptotene spermatocyte nucleoplasm (*) more strongly than large pachytene spermatocytes which display distinct threads. X: SYCP3 stains leptotene spermatocytes (*) which display weak threads and large pachytene spermatocytes with slightly more diffuse threads. XI: SYCP3 stains zygotene spermatocytes (*) with mostly intact threads and large diplotene spermatocytes with slightly diffuse threads. XII: SYCP3 stains zygotene spermatocytes (*) with intact threads and weakly stains M2 spermatocytes with diffuse signal.

**Supp. Table 1. Immunofluorescence Antibodies and conditions**. Antibodies used in this manuscript for immunofluorescence, their species, supplier, product number, and the dilution used.

**Supp. Movie 1. The ADAD2 granule through multiple vertical planes**. ADAD2 granules. Red - SYCP3, green - ADAD2. Slices 0.13 µM apart. 1000x magnification.

**Supp. Movie 2. ADAD2 and RNF17 colocalization across multiple vertical planes**. Red - RNF17, green - ADAD2. Slices 0.13 µM apart. 1000x magnification.

## Notes

### Competing Interest Statement

The authors have declared no competing interest.

